# Comparing Time Series Transcriptome Data Between Plants Using A Network Module Finding Algorithm

**DOI:** 10.1101/564286

**Authors:** Jiyoung Lee, Lenwood S. Heath, Ruth Grene, Song Li

## Abstract

Comparative transcriptome analysis is the comparison of expression patterns between homologous genes in different species. Since most molecular mechanistic studies in plants have been performed in model species including Arabidopsis and rice, comparative transcriptome analysis is particularly important for functional annotation of genes in other plant species. Many biological processes, such as embryo development, are highly conserved between different plant species. The challenge is to establish one-to-one mapping of the developmental stages between two species. In this protocol, we solve this problem by converting the gene expression patterns into a co-expression network and then apply network module-finding algorithms to the cross-species co-expression network. We describe how to perform such analysis using bash scripts for preliminary data processing and R programming language, which implemented simulated annealing method for module finding. We also provide instructions on how to visualize the resulting co-expression networks across species.

## INTRODUCTION

Expression analysis is commonly used to understand the tissue or stress specificity of genes in large gene families [1–5]. The goal of comparative transcriptome analysis is to identify conserved coexpressed genes in two or more species [3,6,7]. The traditional definition of orthologous genes is based solely on sequence homology [8–11] and syntenic relationships [2,12–14] and not on gene expression patterns. In contrast, comparative transcriptome analysis combines a comparison of gene sequences with a comparison of expression patterns between homologous genes in different species. Homologous genes have been reported to be expressed either at different developmental stages, in different tissue types, and/or under different stress conditions [3,15–17]. This documented divergence of expression patterns provides crucial evidence for the existence of functional divergence of homologous genes across species [18,19]. Therefore, comparative transcriptome analysis is an important tool for distinguishing those genes that have retained functional conservation from those that have undergone functional divergence. Comparative transcriptome analysis is particularly important for plant research, since most molecular mechanistic studies in plants have been performed in model species, primarily Arabidopsis [20]. The consequence of this narrow focus is that the functional annotation of the genes of many other plant species relies solely on sequence comparisons with Arabidopsis [21].

To compare transcriptomes between any two species, a first step is to establish homologous relationships between proteins in the two species. A second step is to identify expression data obtained from experiments that are performed under similar conditions or tissue types. The third step is to compare the expression patterns between the two data sets. In this protocol, we will compare published time course seed embryo expression data from Arabidopsis [22] with data from the same tissue in soybean [23] as a demonstration of how to apply computational tools to comparative transcriptome analysis.

In contrast with the time course data examined here, many other datasets have been reported from “treatment-control” experiments (one time point only, two treatment conditions). For example, soybean roots were treated with drought stress in one experiment [4]. To address the question of functional conservation versus functional divergence within gene families, these soybean root data can be compared with transcriptome data from Arabidopsis roots, under a similar stress [24]. This is a relatively simple problem, because, in both experiments, we can identify lists of differentially expressed genes in response to the same or similar treatments. It is a simple two-step process to identify conserved co-expressed genes for treatment-control experiments. First, one needs to identify a list of gene pairs that are homologous between these two species. A simple BLAST search or other more sophisticated approaches such as OMA, EggNog, or Plaza [9,10,12] can be used to identify homologous genes. Second, the two lists of differentially expressed genes can be compared to find whether any pairs of these homologous genes appear in both lists.

In this article, we are focusing on a more complex scenario: two time-series experiments were performed for the same developmental process in two different species [25]. Time course data provide more data points than simple treatment-control experiments and, thus, can reveal relationships based on development between homologous genes in two organisms. However, this is also challenging, because the number of time points in the two experiments are different. It can be challenging to precisely match developmental stages between two species, although some excellent approaches have been proposed [25,26]. Despite the difficulty of establishing one-to-one mapping between the developmental stages of two species, many biological processes, such as embryo development, are known to be highly conserved between different plant species that are compared in comparative transcriptome analysis [27,28]. One way to solve this developmental stage problem is to convert the gene expression patterns into a co-expression network and then apply network alignment or network module-finding algorithms to these co-expression networks [29]. Transforming expression data to a network form simplifies the problem and allows exploration using well established network algorithms [30,31]. In this protocol, we describe how to perform such analysis using a published simulated annealing method [29]. We also discuss how to visualize the resulting co-expression networks across species [32] and the results from different choices of homology finding methods.

### 2. Install software and download experimental data

All scripts used in this analysis can be obtained from github using the following command (Note 4.1).

The “$” means the command is executed under a Linux terminal (Note 4.2).

$ git clone https://github.com/LiLabAtVT/CompareTranscriptome.gitATH_GMA

You can replace “ATH_GMA” with another folder name that better represents your project. All scripts in this project are tested under the project folder created by the “git clone” command (default ATH_GMA).

#### Necessary Resources

This protocol was tested under CentOS 7, which is a Linux operating system. The steps described in this protocol can be used in most UNIX compatible operating systems; this includes all major Linux distributions, and Mac OSX. For Windows users, the individual components of this protocol, such as BLAST, software used for RNA-Seq analysis, programming language R and Python, all have Windows compatible executable files and can be used under Windows environments. In this protocol, we will install NCBI BLAST for the homology search step (Section 2.2), STAR for read mapping and featureCounts for counting reads (Section 2.6), and the R programming language and several packages for RNA-Seq and comparative transcriptome analysis (Section 2.7).

##### 2.1 Set up folder structure for data analysis

To facilitate reproducible and effective computational analysis [33,34], we suggest that the user create a folder structure (**Figure 1**) such that the raw data, processed data, results, and scripts for data processing can be organized into their respective folders. In this protocol, the reader can use the following commands to create the recommended folder structure.

**Figure 1.**
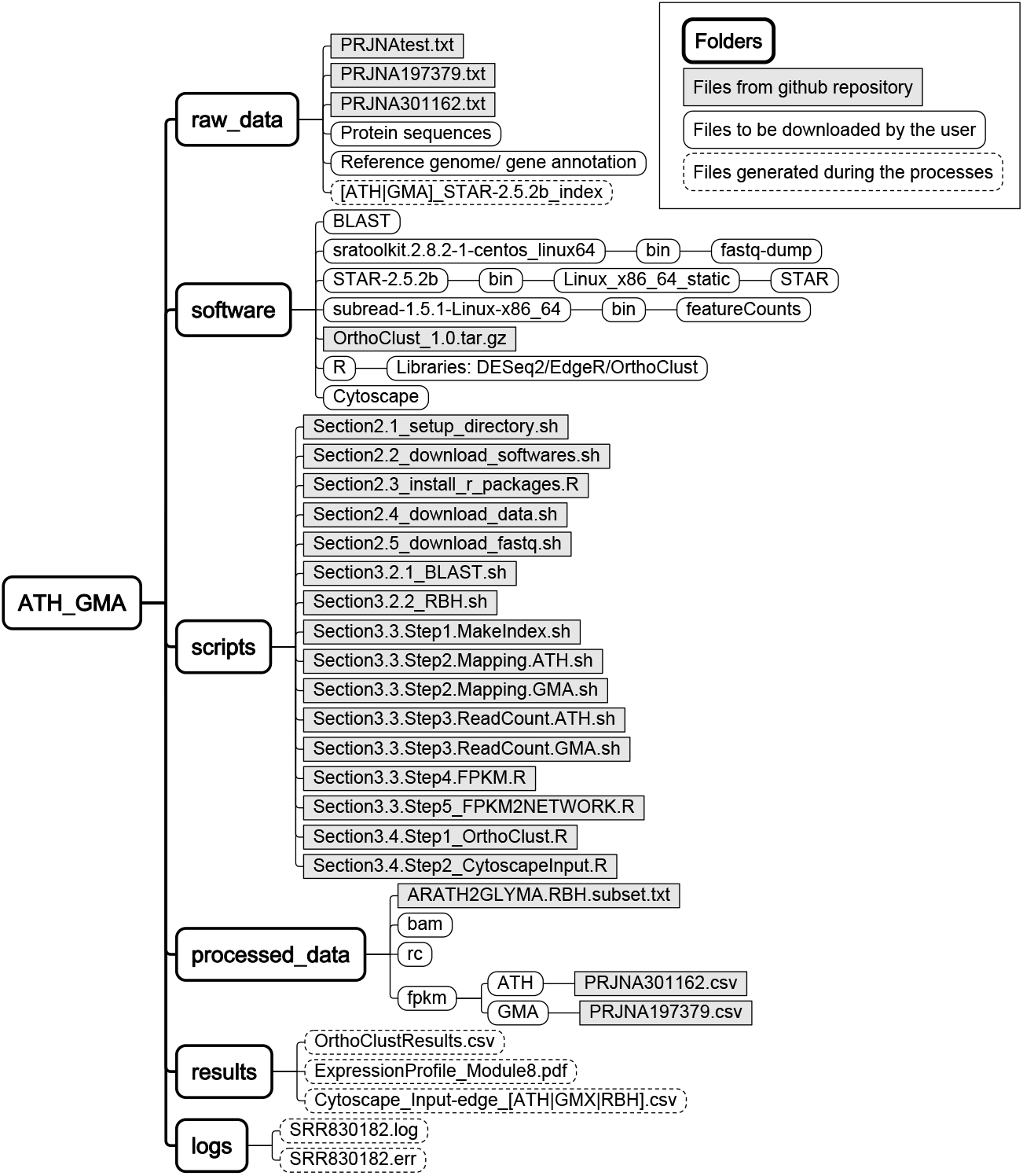
Folder structure for data analysis.

$ cd ATH_GMA
$ mkdir raw_data processed_data scripts results software
$ mkdir processed_data/bam processed_data/rc

Sequence and annotation files from databases should be downloaded to the “**raw_data**” folder. Software tools that will be used in this analysis can be saved and installed in the “**software**” folder. We recommend the reader to create a folder named “**bin**” under the software folder such that the executable files can be copied to “**software/bin**” folder and add “**software/bin**” to the PATH environmental variable under the Linux environment. For experienced Linux users, software can also be installed in a user specified folder such as ~/bin or in a system wide folder. The reader can download scripts in github into the “**scripts**” folder. Intermediate output will be generated in the “**processed_data**” folder, and major input and output files for visualization will be saved in the “**results**” folder.

All scripts for this step are provided in “Section2.1_setup_directory.sh” in the “scripts” folder. The reader can set up the folder structure (**Figure 1**) using the following command.

$ cd ATH_GMA
$ sh./scripts/Section2.1_setup_directory.sh

##### 2.2 Software installation

We provide a script to download and install tools for RNA-seq analysis; readers can run the script in the project folder.

$ cd ATH_GMA
$ sh./scripts/Section2.2_download_softwares.sh

A successfully installed tool will return version information when it is run only with a “–v” or a “— version” option.

###### Install NCBI BLAST for identification of homologous genes

BLAST is a sequence similarity search tool [35]. The latest version of NCBI BLAST can be downloaded from the NCBI ftp site using the following link: ftp://ftp.ncbi.nlm.nih.gov/blast/executables/LATEST/. This folder contains precompiled executable files and installation files for Windows, Mac OSX, and Linux platforms. Because finding orthologous genes at a genome scale is computationally intensive, it is recommended to use a Linux workstation or computing cluster to perform the BLAST analysis.

For Linux users, the current pre-compiled executable is ncbi-blast-2.6.0+-x64-linux.tar.gz.
For Mac users, the current installation file is ncbi-blast-2.6.0+.dmg.
For Windows users, the current installation file is ncbi-blast-2.6.0+-win64.exe.

A later version of BLAST should work as well with minor changes in the command line options. For Windows and Mac users, double click the downloaded file to install the program. For Linux users, one can use “tar-xvf ncbi-blast-2.6.0+-x64-linux.tar.gz” to extract the archive file. After extracting the files, move the executable files to a folder in the Linux search path.

###### Install tools for RNA-Seq data download

The following shows a sample script to download sra-tools and fastq-dump to download the raw sequencing data. The sequence read archive (SRA) database provides sra-toolkit, which is a suite of easy to use computational tools to download data from the database. To download the raw data from the SRA database, one needs to first install the sra-toolkit and use the fastq-dump utility program based on the SRA ids.

$ cd ATH_GMA/software
$ wget http://ftp-trace.ncbi.nlm.nih.gov/sra/sdk/current/sratoolkit current-centos linux64.tar. gz
$ tar-xzf sratoolkit.current-centos_linux64.tar.gz
$ ./sratoolkit.2.8.2-1-centos_linux64/bin/fastq-dump --version

###### Install tools for RNA-Seq data analysis

We will install the STAR [36] and featureCounts [37] software tools. STAR is a read mapper, and featureCounts can count the number of reads mapped to each gene in the genome. Both software tools were used here due to their speed and accuracy [38,39]. Other alternative mappers can be used, and there are excellent review papers [39–41] that compare and summarize these different bioinformatics tools.

To download and install STAR and featureCounts, run the following scripts in the project folder.

$ cd Proj_CompTS_ATH_GMA/software
$ wget https://github.com/alexdobin/STAR/archive/2.5.2b.tar.gz
$ tar-xzf 2.5.2b.tar.gz
$ STAR-2.5.2b/bin/Linux_x86_64_static/STAR-version
$ wget https://sourceforge.net/proiects/subread/files/subread-1.5.1/subread-1.5.1-Linux-x86_64.tar.gz/download
$ tar-zxvf download
$ subread-1.5.1-Linux-x86_64/bin/featureCounts -v

##### 2.3 Install R, DESeq2, and edgeR packages for RNA-Seq data analysis

R is a programing language and environment for statistical data analysis [42]. We will use R to summarize RNA-Seq reads and to generate FPKM data. To install R, the reader should go to the Comprehensive R Archive Network (CRAN) (https://cran.r-project.org) to download the installer packages for their Windows, Mac OSX, or Linux system. For Linux users, R can be installed using the command line, and platform dependent package management systems. For example, to install R in CentOS 7 Linux, the user should simply type:

$ sudo yum install R

Scripts for installing R packages are provided in:

Section2.3_install_r_packages.R

To install DESeq2 and edgeR, the user should follow the instructions for these respective packages. These two packages are part of the Bioconductor repository such that the installation should be performed using the Bioconductor installation script. The following commands are executed under the R environment and these commands are preceded by “>“. For commands that are executed under Linux terminals, these commands are preceded by “$”.

> source(‘https://bioconductor.org/biocLite.R’)
> biocLite(‘DESeq2’)
>biocLite(‘edgeR’)

The installation script will detect the dependency of these two packages and install other required packages accordingly.

To install the OrthoClust package, the user should download the script for the OrthoClust package.

>setwd(“./software”)
>install.packages(‘?rthoClust_1.0.tar.gz”, repos=NULL, type=“source”)

##### 2.4 Download protein and genome sequences for Arabidopsis and soybean

Sample scripts for download are provided in “Section2.4_download_data.sh”. All protein-coding sequences and genomic sequences for Arabidopsis can be downloaded from the Araport web site (www.araport.org). Araport is a data portal for Arabidopsis genomic research that hosts the latest genomic sequences and genome annotations for this model organism [43]. The web site requires free registration to access the download link to the protein sequences and genome annotation files. As of July 2017, the current version of the protein sequences file is “Araport11_genes.201606.pep.fasta.gz”. This name will likely be different for future versions of the protein sequences. We recommend that users download the latest version of the protein sequences, and record the actual download date and version of the sequence files for the purpose of reproducibility. The latest version of the genome sequence of Arabidopsis is “TAIR10_Chr.all.fasta.gz”. This file is unlikely to change because the genome assembly of Arabidopsis is likely to remain the same in the future. The latest version of the gene annotation file is “Araport11_GFF3_genes_transposons.201606.gtf.gz”.

All protein-coding sequences for soybeans can be downloaded from the DOE phytozome database (https://phytozome.jgi.doe.gov/pz/portal.html#!bulk?org=Org_Gmax). Phytozome is a data portal for plant and microbial genomes that hosts dozens of sequenced plant genomes and gene annotations [44]. This web site also requires free registration before data downloading. The latest version of soybean protein sequences is version 2.0 (downloaded in July 2017). The protein sequences and genomic sequences are “Gmax_275_Wm82.a2.v1.protein.fa.gz” and “Gmax_275_v2.0.fa.gz”. These names are likely to change with future versions of the genome and proteome annotation. The latest version of the gene annotation file is “Gmax_275_Wm82.a2.v1.gene_exons.gff3.gz”.

These files are in compressed fasta format and require de-compression before use. Under the Linux command line, the following command can be used to de-compress these “*.gz”.

$ gunzip Araport11_genes.201606.pep.fasta.gz
$ gunzip Gmax_275_Wm82.a2.v1.protein.fa.gz

##### 2.5 Download raw data from published RNA-Seq experiments

Raw sequencing data can be downloaded from the NCBI Sequence Read Archive (SRA) (https://www.ncbi.nlm.nih.gov/sra). The embryo developmental data sets for Arabidopsis and soybean can be found in two bioprojects (PRJNA301162 for Arabidopsis and PRJNA197379 for soybean). For the Arabidopsis samples, RNA-Seq data were collected in triplicates at seven time points (7, 8, 10, 12, 13, 15, and 17 days after pollination). For the soybean samples, RNA-Seq data were collected in triplicates at ten time points (5, 10, 15, 20, 25, 30, 35, 40, 45, and 55 days, day 0 of the time course is 12 to 17 days after anthesis). Each sample is represented by a unique GSM id; for example, the three replicates of 7 days old Arabidopsis embryo samples are GSM1930276, GSM1930277, and GSM1930278. All 41 samples from this experiment are stored under a unique GSE id, GSE74692. Each sample is also represented by a unique SRA id. For example, the three replicates of 7 days old Arabidopsis embryo samples are SRR2927328, SRR2927329, and SRR2927330 from PRJNA301162.

$ fastq-dump--split-3 SRR2927328-outdir./raw_data

We suggest that the reader download the data into the raw data folder for further processing. To download large numbers of data sets, prepare a text file with all SRR ids for one species and run the following script in the project folder.

$ cd ATH_GMA
$ sh./scripts/Section2.5_download_fastq.sh./raw_data/PRJNA301162.txt ATH
$ sh./scripts/Section2.5_download_fastq.sh./raw_data/PRJNA197379.txt GMA

Depending on the size of sequencing data and network speed, this step may take a few hours. We provide a test file “PRJNAtest.txt” for the user to test the execution time for downloading one file. The time for downloading the entire data set can be estimated based on downloading this single file. We also provide the FPKM data for this particular data set so that the users do not need to download the original data to perform the analysis in this protocol. To perform the analysis using provided FPKM file, the user can start the analysis from Section 3.4.

## 3 Methods

### 3.1 Comparative transcriptome analysis overview

This protocol provides details of comparative transcriptome analysis between two species. We not only compute sequence similarity between protein coding genes in two species, we also integrate the gene expression patterns of these genes from two different species under similar biological processes. There are three major steps in this analysis (**Figure 2**): 1) identify homologous genes between two species; 2) generate a gene expression data matrix and a co-expression network in each species; 3) perform cross species comparisons of gene homology and expression patterns. For each of these steps, multiple bioinformatics tools are available. This protocol will provide a basic workflow for each of the steps and the reader can substitute individual steps with other tools (**See Note 4.3**).

**Figure 2.**
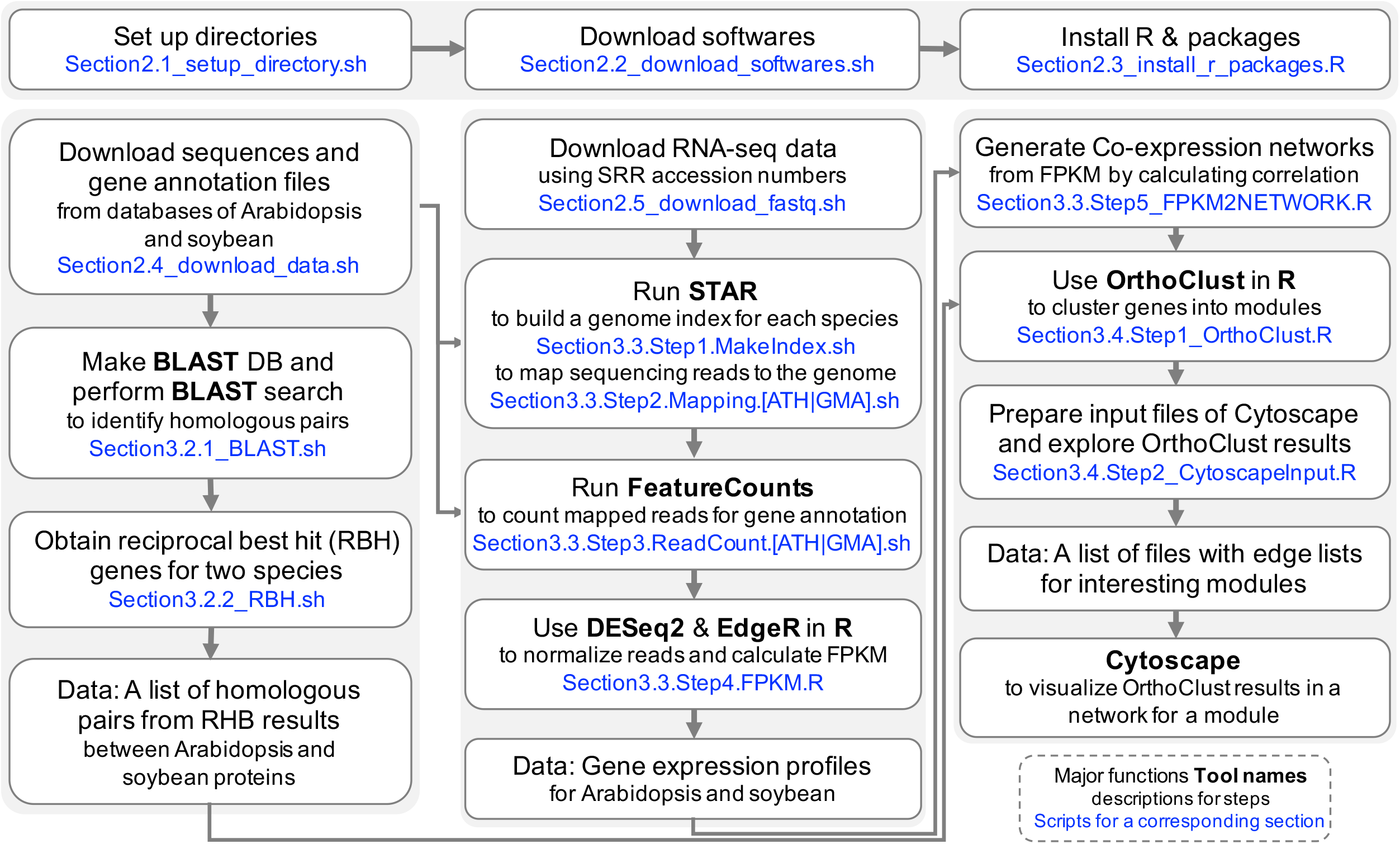
A workflow of comparative transcriptome analysis between soybean and Arabidopsis. It is composed of three major parts: identification of ortholous pairs between two species using BLAST, RNA-seq analysis to get co-expression networks, and running OrthoClust to cluster genes with orthologous relations. Blue fonts indicates softwares or scripts used in this workflow.

### 3.2 Identifying homologous genes between species

#### 3.2.1 Identification of homologous pairs using BLAST

Analysis in this section can be performed using the following command:

$ cd ATH_GMA
$ sh./scripts/Section3.2.1_BLAST.sh

**Step 1.** Merge the Arabidopsis protein fasta file and soybean protein fasta file using this Linux command:

$ cat Araportii.pep.fasta GLYMA2.pep.fasta > ATHGMA.pep.fasta

**Step 2.** Create the BLAST database:

$ makeblastdb-in ATHGMA.pep.fasta \
-out ATHGMA.blastdb \
-dbtype prot \
-logfile makeblastdb.log

The option “–in” specifies the input file name of the merged protein fasta file. “–out” specifies the BLAST database file name. “–dbtype” indicates the database is a protein database. “–logfile” is for recording error messages in case the process fails.

**Step 3.** Perform the BLAST search.

The Linux command used in this step is:

$ blastp-evalue 0.00001 \
-outfmt 6-db ATHGMAX.blastdb \
-query ATHGMA.fasta > ATHGMA.pep.blastout

The option “–evalue” specifies the E value threshold. “–outfmt” is set to be 6, which is tab delimited format. “–db” is set to be the BLAST database built in step 3. “–query” uses the merged protein fasta files as input. The results of BLAST analysis are written in a file named ATHGMAX.pep.blastout.

The output includes the following 12 tab-separated columns “**qseqid sseqid pident length mismatch gapopen qstart qend sstart send evalue bitscore**”. The meaning of these columns can be found using the BLAST help manual. The columns that will be used in downstream analysis are **qseqid** (query sequence id), **sseqid** (subject sequence id), and **evalue** (E value). We will filter BLAST results and only keep homologous genes with BLAST E value < 1e-5 [3,26].

#### 3.2.2 Obtaining reciprocal best hit (RBH) genes

Reciprocal best BLAST hit (RBH) and its variants are commonly used methods to identify homologous genes in two species [45–49]. To identify RBH genes between any two species, the BLAST results from protein sequence alignment were first parsed to identify the best BLAST hit for each soybean protein in the Arabidopsis protein lists. For each soybean protein, there is at most one best BLAST hit protein in the Arabidopsis proteome. For each of the Arabidopsis proteins identified in the first step, the best BLAST hit of each protein in the soybean proteome is also identified. If this best hit is also the original homologous gene found in the first step, this pair of proteins is defined to constitute an RBH pair.

For genes with multiple isoforms and potentially multiple protein sequences, we performed the BLAST analysis at the isoform level and then collapsed all the isoforms for each gene to find the best match. In fact, a large fraction of the isoforms in both Arabidopsis and soybean do not change their protein coding sequences, the difference being found in the UTR regions of the transcripts being compared. This is consistent with published results in Arabidopsis and soybean [50,51]. We developed a Python script that can identify RBH genes from the above two species from BLAST results. The user can download this script from the github repository. To perform the analysis the user can use the following commands:

$ cd ATH GMA
$sh./scripts/Section3.2.2_RBH.sh

Although RBH genes are widely used in comparative genomic analysis, other methods can be used to identify homologous genes for downstream analysis (see **Note 4.3**). An example file (ARATH2GLYMA.RBH.subset.txt) of RBH genes is provided. The user can use this file to perform the following analysis without running the RBH script.

### 3.3 Gene expression data processing

Gene expression quantification includes three main steps: 1) read mapping; 2) read counting and 3) FPKM calculation. For this analysis, we follow a published protocol for expression processing [50].

**Step 1**. Create genome index by STAR.

RNA-Seq reads have to be mapped to the respective reference genomes. To use STAR to map reads to the reference genome, the user needs to build a genome index using the following commands.

$ cd ATH_GMA
$ sh./scripts/Section3.3.Step1.MakeIndex.sh

The following commands are used to create a genome index for Arabidopsis.

$ WORKDIR=$(pwd)
$ IDX=$WORKDIR/raw_data/ATH_ST AR-2.5.2b_index
$ GNM=$WORKDIR/raw_data/T AIR 10_Chr.all.fasta
$ GTF=$W0RKDIR/raw_data/Araport11_GFF3_genes_transposons.201606.gtf
$ STAR --runMode genomeGenerate \
--genomeDir $IDX \
--genomeFastaFiles $GNM \
--sjdbGTFfile $GTF

The option “–-runMode” indicates that the command is to create a genomic index. “–-genomeDir” specifies the file name for the genome index. --genomeFastaFiles” indicates the input fasta file for genomic sequences. “–-sjdbGTFfile” is to provide a genome annotation file when creating the genomic index. A genome index will be created for each species.

**Step 2.** Read mapping by STAR.

After creating genome indexes, the user needs to use STAR to map reads from each sample to the reference genome to generate a read mapping file using the following commands.

$ cd ATH_GMA
$ sh./scripts/Section3.3.Step2.Mapping.ATH.sh
$ sh./scripts/Section3.3.Step2.Mapping.GMA.sh

The “Section3.3.Step2.Mapping.ATH.sh” is to map all Arabidopsis reads. The “Section3.3.Step2.Mapping.GMA.sh” is to map all Soybean reads. In the SRA database, each sample has a unique SRR id. The following commands show one example of such SRR ids (SRR2927328). SRR2927328_1 and SRR2927328_2 represent two ends of paired reads.

$ STAR --genomeDir $IDX \
--readFilesIn $WORKDIR/raw_data/SRR2927328_1.fastq.gz
$WORKDIR/raw_data/SRR2927328_2.fastq.gz \
--outFileNamePrefix $WORKDIR/processed_data/bam/SRR2927328/SRR2927328 \
--outSAMtype BAM SortedByCoordinate

The option “–-genomeDir” specifies the file name for the genome index. “–-readFilesIn” indicates the input fastq files for RNA-seq reads. Two files are provided for paired-end reads. --outFileNamePrefix” is to provide the directory for output data. “—outSAMtype BAM” indicate the output file should be a bam file. “SortedByCoordinate” set the output data to be sorted by the order of where the read is mapped to the chromosome.

**Step 3**. Read counting with featureCounts.

To count reads with featureCounts, the user can use the following command:

$ cd ATH_GMA
$ sh ./scripts/Section3.3.Step3.ReadCount.ATH.sh
$ sh ./scripts/Section3.3.Step3.ReadCount.GMA.sh

For this step, featureCounts will calculate how many reads map to each gene region. For simplicity, we only count uniquely mapped reads and only summarize read counts at the gene level. Other software can be used to summarize expression at isoforms levels. The following commands are for counting reads for a single file.

$ WORKDIR=$(pwd)
$ GTF=$W0RKDIR/raw_data/Araport11_GFF3_genes_transposons.201606.gtf
$ BAM=$WORKDIR/processed_data/bam
$ RC=$WORKDIR/processed_data/rc
$ featureCounts -t exon \
-g gene_id \
-p \
-a $GTF \
-o $RC/SRR2927328.readcount.txt \
$BAM/SRR2927328/SRR2927328Aligned.sortedByCoord.out.bam

The option “–t exon” indicates that only reads mapped to exons are counted. The option “–p” indicate the input reads are paired-end reads. The option “–a” provides the location of the genome annotation file. The option “–o” specifies the output file location. The last parameter is the file name of the read mapping file (bam file).

**Step 4.** FPKM calculation using DESeq2 and edgeR.

For this step, R scripts will be used to summarize gene expression level in fragments per kilo-basepairs per million reads (FPKM). To calculate FPKM, we performed the following five steps: 1) merging read counts from different files into one single file; 2) differential expression analysis using DESeq2; 3) data normalization. 4) FPKM calculation and 5) average FPKM calculation across replicates. These steps can be performed using a unified sh (shell) script: NGS_RNA-seq_CalcFPKM.R, which is provided in the github repository of this project. To run this script, the user needs to provide a table that summarizes the replicate structure of the samples. Example tables (PRJNA301162.csv for Arabidopsis and PRJNA197379.csv for soybean) are provided in the “processed_data” folder.

To run the unified R script for FPKM calculation, use the following commands:

$ cd ATH_GMA
$ Rscript ./scripts/Section3.3.Step4.FPKM.R ./processed_data/fpkm/GMA
$ Rscript ./scripts/Section3.3.Step4.FPKM.R ./processed_data/fpkm/ATH

This script requires multiple input files to be present in the working directory. These files include a file that describes the design matrix of the experiment and the read count files generated in Step 3. More descriptions of the input file formats are included in the annotation of the R script.

**Step 5**. Co-expression Networks from gene expression profiles

Expression data will be summarized and converted to gene co-expression networks. The input data include data matrices with averaged and normalized FPKM values. In this protocol, we use genes in metabolic pathways that are essential to seed development. Other methods can be used to filter genes before the analysis, for example, only keep genes with high variations across conditions. Finally, gene coexpression matrices were calculated for each species. We use the cut-off with p value < 0.001 and Pearson Correlation Coefficient > 0.99 to generate co-expression networks. To generate co-expression networks, the following commands were used.

$ cd ATH_GMA
$ Rscript ./scripts/Section3.3.Step5_FPKM2NETWORK.R

### 3.4 Identify orthologous co-expressed clusters using OrthoClust

#### 3.4.1 Overview of the OrthoClust method

Simple approaches can be used to identify conserved co-expression genes across different species. For example, one can first cluster gene expression in two species separately, and, for each pair of cluster combinations, one can find whether the pairs of clusters share significantly large numbers of homologous genes using appropriate statistical tests such as Fisher’s exact test. OrthoClust [29] is a global approach in which the process of co-expression clustering finding and homology detection is integrated into the same objective function. The objective function *H* is defined as

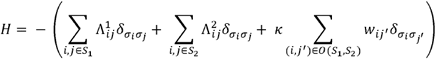

where *S_N_* is the sets of genes for a species and a subscript of *S* (N = 1 or 2) corresponds to the species respectively. *i* and *j* are individual genes of a species or nodes on a network. 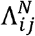 denotes a modularity score from gene *i* and *j*, that is a difference between the real number of edges and the expected number of edges. *δ_σ_i_σ_j__* is for a module label. If *i* and *j* have the same module label, *δ_σ_i_σ_j__* =1, and, if not, *δ_σ_i_σ_j__* =0. A coupling constant, *κ* controls overall impact of orthology relations on the objective function, and a weight, *W_i,j_* is for orthology relations coming from the number of orthologus genes between two species. The objective function Hwill return lower values when orthologous genes are assigned into the same module.

This approach translates orthologous co-expression finding into a network module finding problem. The objective function includes three components: two components represent the goodness of the expression clustering results and one component represents the effect of homologous genes across species. The parameter *κ* can be adjusted to increase or decrease the contribution of homologous genes in the clustering processes. The effects of using different co-expression thresholds and parameter *κ* are discussed in Note 4.4.

#### 3.4.2 Steps for OrthoClust analysis

To perform OthoClust analysis, we require three input data files: 1) the gene co-expression network from soybean; 2) the gene co-expression network from Arabidopsis; and 3) the orthologous gene pairs between two species.

These files require a specific format for the OrthoClust engine to analyze. The user can use the following R command to perform the clustering analysis

> library(OrthoClust)
>OrthoClust2(Eg1=GMX_edgelist, Eg2=ATH_edgelist, \
list_orthologs=GA_orthologs, kappa=3)

We provide a wrapper script that will read three input files: a list of edges from Arabidopsis, a list of edges from soybean, and a list of RBH gene pairs from two species. To perform OrthoClust analysis, the user can simple use the following commands:

$ cd ATH_GMA
$ Rscript ./scripts/Section3.4.Step1_OrthoClust.R

This script will generate three files. “Orthoclust_Results.csv” contains information regarding modules assignment for each gene. “Orthoclust_Results_Summary.csv” includes number of genes assigned to each module. “Orthoclust_Results.RData” contains multiple R objects that will be used in the visualization step.

#### 3.4.3 Visualization of OrthoClust results as a network

To visualize OrthoClust results, we use Cytoscape, a network visualization platform to analyze biological networks and to integrate multiple data into networks such as gene expression profiles or annotation [52]. We used module 8 from the previous step as an example. There are three input files: 1) soybean coexpression network edge list for genes in module 8, 2) Arabidopsis co-expression network edge list for genes in module 8, and 3) RBH list for genes in module 8. To generate these files for Cytoscape visualization, the user can use the following command.

$ cd ATH_GMA
$ Rscript ./scripts/Section3.4.Step2_CytoscapeInput.R

**Step 1**. To Import three files on Network Browser, we can first start from the Cytoscape menu bar “File” > “import” > “Network” > “File”. After you select one of three input files, the popup window with “Import Network From Table” title appears. You can see two columns with gene names in the middle of the window. Next, to change attributes of columns, click the first line of each column and choose either “Source Node” or “Target Node” from the menu. Since three edge lists do not have direction, the two columns from each input file can be assigned into either source or target nodes. After that, we change an option for column names from “Advanced Options” at the bottom left of the window. On the new popup window, we can uncheck “Use first line as column names”, since we do not have headers in the input files. Finally, you can see two column names, “Column1” and “Column2” with different icons of attributes, and the remaining parts of the preview are gene names. You can repeat these steps for each of the input flies.

**Step 2**. With three imported networks, we can integrate data sets of co-expression networks with homologous relations using the Union function. To do that, select three network on the network tab on the control panel (click one network and click the other two networks while pressing Command), and move to Cytoscape”s menu bar “Tools” > “Merge” > “Networks”.

In the popup window for “Advanced Network Merge”, we should choose the “Union” button, select three networks from “Available Networks”, and then click the right-facing arrow acting for “Add Selected”. After that you can find that three networks are now on “Networks to Merge”, and you can click “Merge” button to merge three networks.

The name of the merged network will appear with the total number of merged nodes and edges on the Network tab on the control, and usually it is automatically visualized on the Cytoscape canvas.

**Step 3**. To express properties of networks (species information, source of edges such as co-expression networks or homologous relations), we can customize visual attributes of the merged network. To do that, on the Select tab on the control panel, we can click the “+” icon below the “Default filter” and choose “Column Filter” to add the new condition. From the “Choose column” drop-down list, you can select “Node: name” or “Edge: name” and type a prefix of each species (“AT” for Arabidopsis genes, or “Glyma” for soybean genes). This filter applies to visualization of the merged automatically, so you can see highlighted nodes on the Cytoscape canvas.

There are several ways to change visualization properties of the selected components. First, we can set “Bypass Style” for the selected nodes or edges such as “Fill Color” and “Size” for properties of nodes, or “Stroke Color” and “Line Type” for properties of edges. To do this, move your mouse pointer on one of the highlighted nodes, right-click, and then select “Edit” > “Bypass Style” > “Set Bypass to Selected Nodes” on the popup menu. The control panel on the left side will be automatically changed to the “Style” tab, and you can see three subtabs: “Node”, “Edge”, and “Network” on the bottom of the interface. Second, we can apply different Layouts with these selected nodes or all nodes from Cytoscape menu bar “Layout”.

As an example of the network with module 8, nodes and edges from soybean and Arabidopsis genes were switched to green and orange colors respectively. To highlight genes of interest, we used thicker double lines for edges and blue color for nodes. We separated genes into four groups according to their input files and species (Arabidopsis genes from RBH results or not, and soybean genes from RBH results or not), and layout each of them with Degree Sorted Circle Layout (Figure 3).

**Figure 3.**
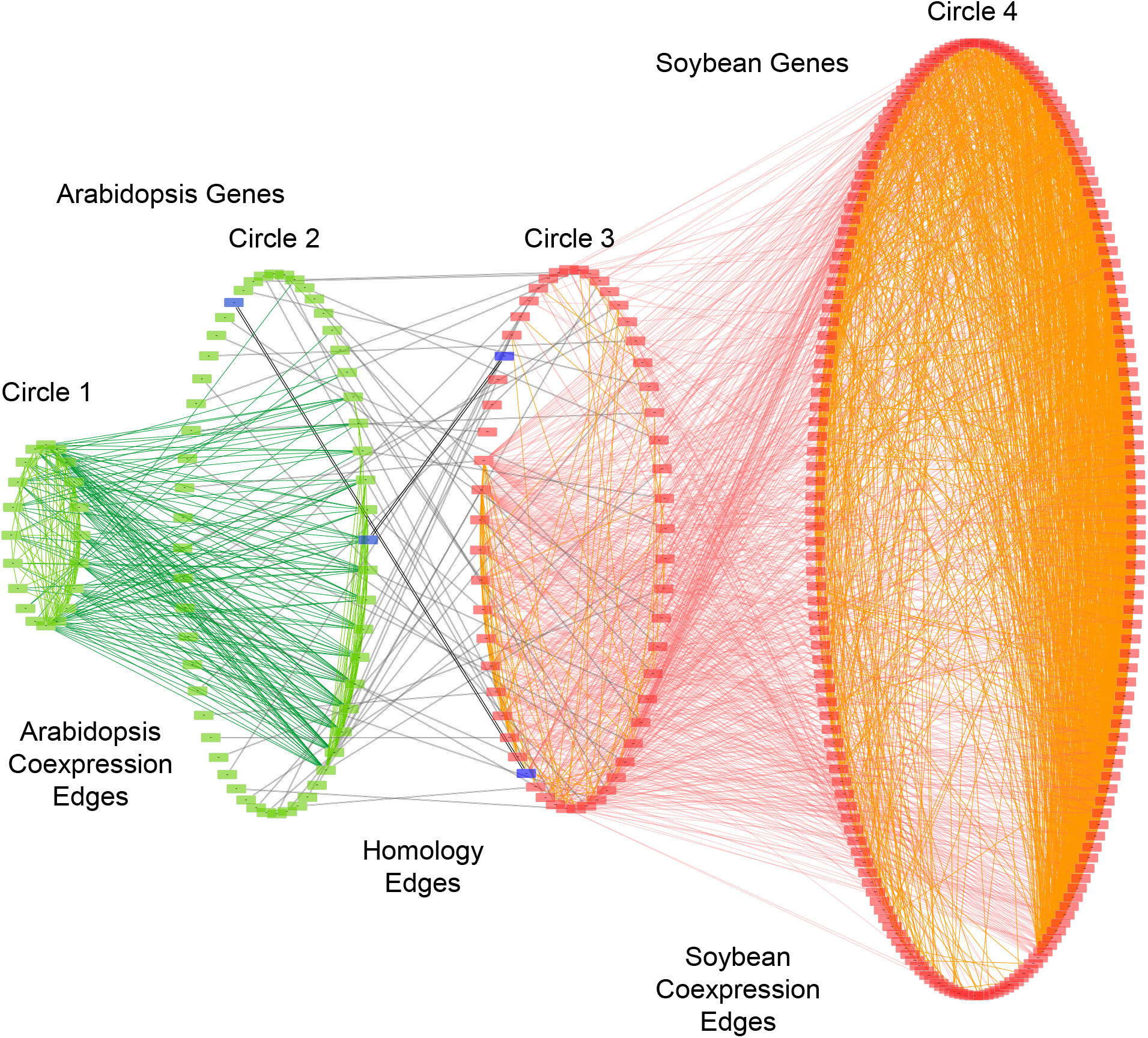
Visualization of module 8 from OrthoClust result. In this network, Circle 1 and 4 stand for groups of genes from Arabidopsis and soybeans that do not have orthology in the other species and only co-expression partner from the same species. Circle 2 and 3 denote genes have orthologous partner in the other species as well as their co-expression partners from the same species. Green nodes are genes from Arabidopsis, and red from soybean. Edges from co-expression network of Arabidopsis are green, and those of soybeans are red. Black double lined edges indicate homologous pairs between soybean and Arabidopsis genes. Four genes from raffinose biosynthesis pathways are highlighted in blue color and their homologous pairs have thicker edges.

#### 3.4.4 Visualization of OrthoClust results as expression profiles

We also provide scripts to directly visualize gene expression patterns for orthologous co-expression modules (**Figure 4**). This figure is generated by the script “Section3.4.Step2_CytoscapeInput.R”. In this module, most soybean genes are tightly clustered. Some Arabidopsis genes are tightly clustered (close to the black line) whereas other Arabidopsis genes are not. This result shows that many genes in the soybean co-expression cluster change their expression patterns in Arabidopsis, suggesting potential functional divergence of these genes. In contrast, many genes that are RBH pairs in the two species have similar expression patterns. For example, one gene (AT5G52560, green line) that is related to the raffinose biosynthetic pathway has a similar decreasing expression pattern as its RBH gene (Glyma.04G245100) in soybean.

**Figure 4.**
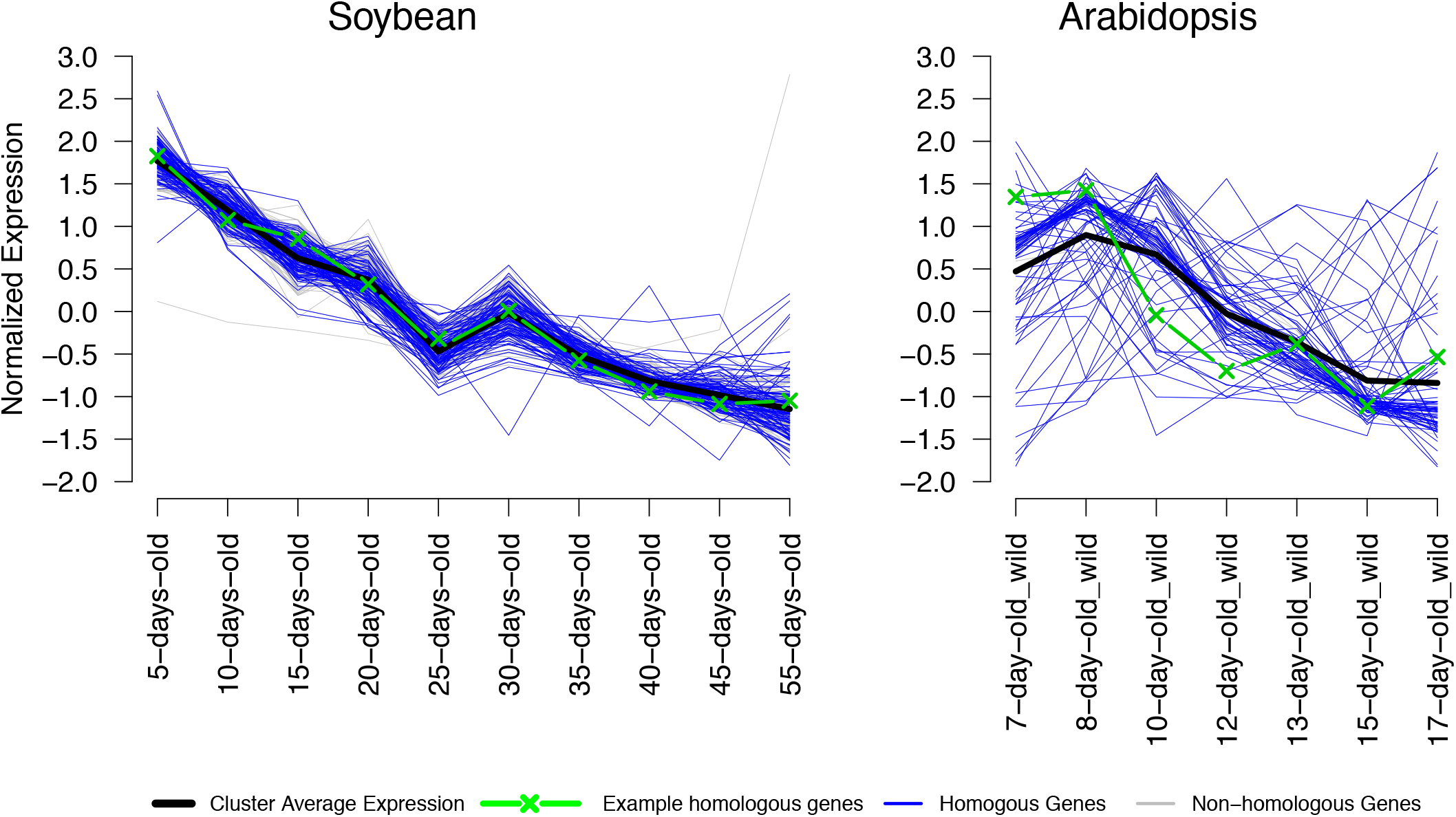
Expression plots of genes from Arabidopsis and soybean bellowing to one of modules of OrthoClust result. One example of homologous genes in Arabidopsis and soybeans are AT5G52560 and Glyma.04G245100 are highlighted in green.

## 4. Notes

### 4.1 Software installation

The git software is installed in most Linux systems by default. If git is not installed in your system, please refer to https://git-scm.com for installation instructions.

### 4.2 Code blocks

All code blocks started with “$” are command line scripts that should be executed under a Linux terminal. All code blocks started with “>“ are command line scripts that should be executed under an interactive R programming language console.

### 4.3 For each of these steps, multiple bioinformatics tools are available

This protocol will provide a basic workflow for each of the steps and the reader can substitute individual steps with other tools. For example, in searching for homologous genes, several other alternative tools such as OMA or OrthoFinder [10,11] can be used instead of BLAST. A comprehensive comparison of these tools is out of the scope of this chapter. Some databases or tools provide pre-computed homologous genes [8,12]. Additional steps must be performed to ensure that the gene ids from OMA, OrthoFinder, or PLAZA match the gene ids used in the expression analysis.

### 4.4 Many genes in both species were not included in the RBH gene lists

This is because the criterion for identifying RBH genes is highly stringent, as it requires that both genes in two species be the best BLAST hit in their respective species. This can be relaxed to identify k-best-hits in two species [6]. We have developed a script that can generate k-best-hits using BLAST results between any two species (OrthologousGenes_OneWayTopNBestHit.py).

### 4.5 Effect of different parameters in OrthoClust analysis

We analyzed how different parameters affect the results of this analysis. We focus on two major parameters (**Figure 5**): the Pearson Correlation Coefficient (PCC) threshold that was used to convert co-expression data to networks, and the kappa parameter that was used in OrthoClust analysis.

**Figure 5.**
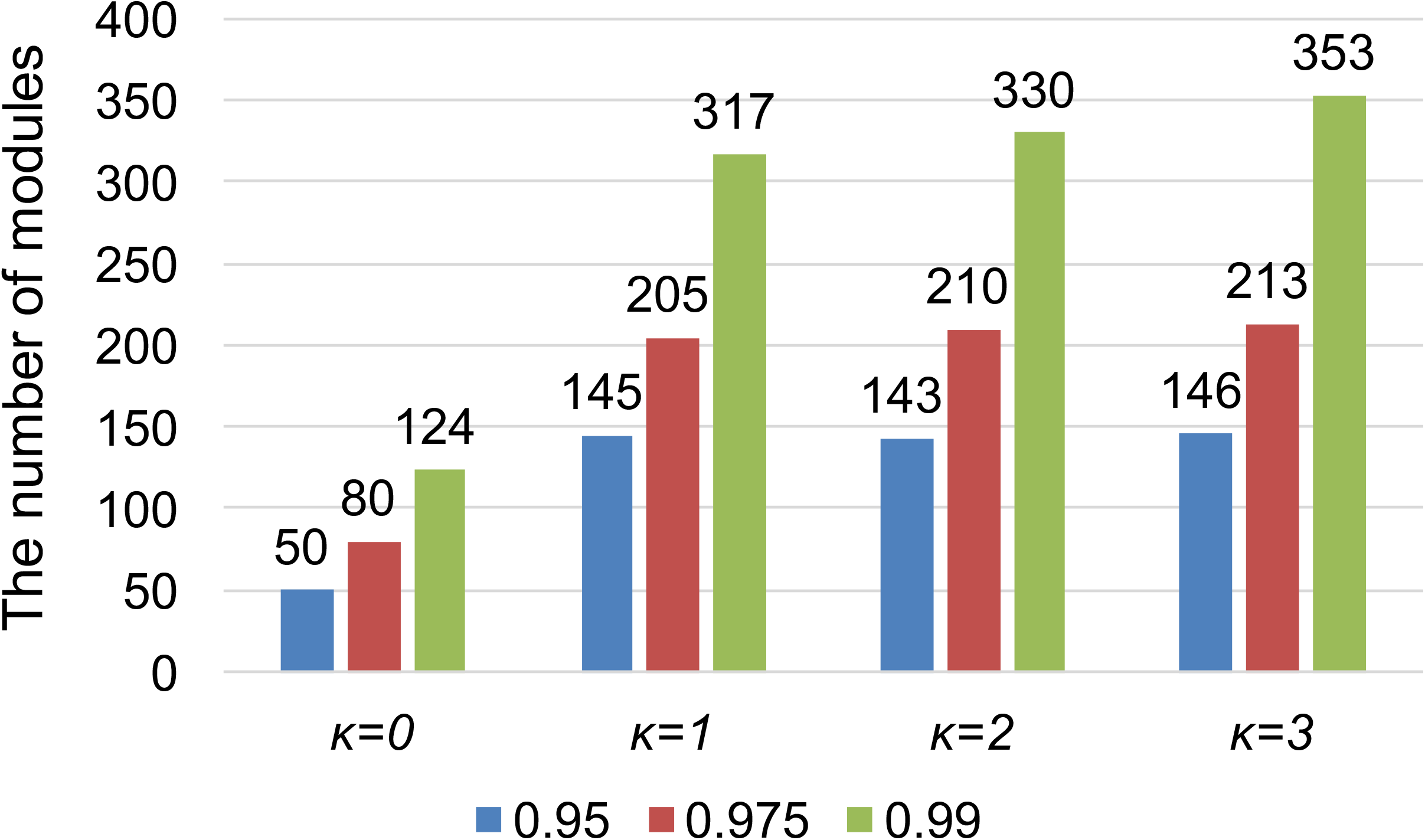
Effect of different correlation cutoff and *κ* values on the number of modules in orthoClust analysis.

The kappa parameter is used to adjust the relative importance of the co-expression edges and homologous edges in network module finding algorithms. When kappa equals zero, the module finding method only finds co-expression modules and does not consider the effects of homologous edges. When kappa is set to be higher than zero, homologous edges will be included in the module finding objective function. This can be verified by comparing the numbers of modules found by kappa = 0 to numbers of modules found by kappa > 0. The numbers of modules found by kappa = 1 is 2 to 3 times the numbers of modules found by kappa = 0. This result suggests that including homologous edges generates more modules across species, because, when kappa = 0, all modules are from the same species. Comparing the numbers of modules from kappa = 2 with kappa = 1, and kappa = 3 with kappa = 2 suggest that increasing kappa can further increase the number of modules.

The PCC threshold also affects the number of modules identified. For the same kappa value, a higher PCC threshold always leads to more modules. This is expected as a co-expression network with higher PCC threshold contains fewer edges. Because of the reduced number of edges, the network is less connected and can be break into more modules as compared to the network generated with lower PCC threshold.

## Ethics approval and consent to participate

Not applicable

## Consent for publication

Not applicable

## Availability of data and materials

The datasets and software supporting the conclusions of this article are available in the Github repository (https://github.com/LiLabAtVT/CompareTranscriptome).

## Authors’ contributions

JL and SL designed the analysis. RG provided the original data and interpreted the biological results. LH edited the manuscript and provided suggestions to improve the methods. JL developed the methods. SL and JL wrote the manuscript.

## Acknowledgments and Funding

This work is partly supported by Virginia Soybean Board.

## Key Reference

Yan et al., 2014. See above.

A methodology to cluster integrated data from co-expression profile for each species and from homologous relationships between multiple species.

## Internet Resources

https://www.araport.org

*Arabidopsis information portal*

https://phytozome.jgi.doe.gov/pz/portal.html#!bulk?org=Org_Gmax

*Genomics resource page of Glycine max Wm82.a2.v1in Phytozome*

https://git-scm.com

*Git software Home page*

https://github.com/LiLabAtVT/CompareTranscriptome.git

*Github page for this tutorial*

https://www.ncbi.nlm.nih.gov/sra

*NCBI Sequence Read Archive (SRA) Home page*

https://github.com/alexdobin/STAR

*Github page of STAR*

http://bioinf.wehi.edu.au/subread-package/

*The Subread package Web page*

https://cran.r-project.org

*The Comprehensive R Archive Network Web page*

**Table 1.** Results of Identified Orthologous Genes.

| Species | Soybean | Arabidopsis |
| --- | --- | --- |
| **Number of proteins** | 48,375 | 24,148 |
| **(Total number of gene models)** | (56,044) | (37,336) |
| **Blast results in each species** | 1,086,080 | 1,081,623 |
| **(Query: Blast DB)** | (Soybean: Arabidopsis) | (Arabidopsis: Soybean) |
| **Number of RBH genes in each species** | 13,024 | 13,024 |
| **Number of 5 best hit in each species** | 208,343 | 112,819 |

**Table 2.** Examples of input data files for OrthoClust analysis. There are three inputs: two co-expression networks of (A) soybean and (B) Arabidopsis, (C) orthologous pairs between soybean and Arabidopsis.

| (A) |  | (B) |  | (C) |  |
| --- | --- | --- | --- | --- | --- |
| row | column | row | column | Soybean gene | Arabidopsis gene |
| Glyma.01G006400 | Glyma.01G016500 | AT1G01540 | AT1G05350 | Glyma.01G001300 | AT2G07050 |
| Glyma.01G021300 | Glyma.01G021400 | AT1G06040 | AT1G06150 | Glyma.01G005800 | AT4G29310 |
| Glyma.01G019400 | Glyma.01G022500 | AT1G01720 | AT1G07400 | Glyma.01G006100 | AT4G26300 |
| Glyma.01G015400 | Glyma.01G026700 | AT1G05230 | AT1G07570 | Glyma.01G010100 | AT1G32090 |
| Glyma.01G025100 | Glyma.01G026700 | AT1G02660 | AT1G08230 | Glyma.01G015400 | AT2G35470 |
| Glyma.01G025100 | Glyma.01G028900 | AT1G01090 | AT1G08510 | Glyma.01G019400 | AT5G65670 |

**Table 3.** Top 10 OrthoClust results sorted by the total number of genes from a module. OrthoClust was performed with parameters ĸ=3, gene co-expression correlation cutofÊĩcorrelaand homologous pairs obtained from RBH Blast.

| No. | Module ID | Total number of genes from a module | The number of genes from soybean | The number of genes from Arabidopsis |
| --- | --- | --- | --- | --- |
| 1 | 2 | 352 | 273 (77.6%) | 79 (22.4%) |
| 2 | 8 | 331 | 255 (77.0%) | 76 (23.0%) |
| 3 | 39 | 297 | 174 (58.6%) | 123 (41.4%) |
| 4 | 1 | 253 | 214 (84.6%) | 39 (15.4%) |
| 5 | 3 | 245 | 207 (84.5%) | 38 (15.5%) |
| 6 | 187 | 215 | 56 (26.0%) | 159 (74.0%) |
| 7 | 224 | 212 | 53 (25.0%) | 159 (75.0%) |
| 8 | 57 | 192 | 110 (57.3%) | 82 (42.7%) |
| 9 | 113 | 147 | 38 (25.9%) | 109 (74.1%) |
| 10 | 19 | 45 | 39 (86.7%) | 6 (13.3%) |

